# Deletion disrupts a conserved antibody epitope in a SARS-CoV-2 variant of concern

**DOI:** 10.1101/2021.03.05.434168

**Authors:** Linda J. Rennick, Lindsey R. Robinson-McCarthy, Sham Nambulli, W. Paul Duprex, Kevin R. McCarthy

**Affiliations:** The Center for Vaccine Research, University of Pittsburgh School of Medicine, Pittsburgh PA; The Department of Microbiology and Molecular Genetics, University of Pittsburgh School of Medicine, Pittsburgh PA; Hillman Cancer Center, Pittsburgh PA; The Department of Pathology, University of Pittsburgh School of Medicine, Pittsburgh PA

## Abstract

Multiple SARS-CoV-2 variants with altered antigenicity have emerged and spread internationally. In one lineage of global concern, we identify a transmitted variant with a deletion in its receptor binding domain (RBD) that disrupts an epitope which is conserved across sarbecoviruses. Overcoming antigenic variation by selectively focusing immune pressure on this conserved site may, ultimately, drive viral resistance.

## Main Text

The SARS-CoV-2 spike (S) glycoprotein is the target of protective antibodies and sole antigen delivered in widely deployed vaccines ^1-4^. Emergent variants (B.1.1.7, B.1.351, P.1) have independently acquired a common series of mutations that confer resistance to therapeutic antibodies and serum from infected or vaccinated individuals. Variant spread has been rapid. Reformulated therapies and vaccines are in development. Directing antibodies to conserved sites therapeutically, as a consequence of immune recall or via designed immunogens can, in theory, overcome antigenic variation. These strategies rely upon sites that slowly, if ever, accrue diversity. We identify a transmitted B.1.1.7 variant with a deletion in a site of pan-sarbecoviruses conservation. This deletion disrupts the binding of an antibody that engages both SARS-CoV and SARS-CoV-2. The acquisition of antigenic novelty in S glycoprotein has been recurrent and convergent. By extension conservation at this site may not persist.

Using sequences deposited in the GISAID database^5^, we have monitored variants of concern for the acquisition of additional epitope-altering mutations. We identified a transmission cluster of six identifiably different individuals that share a nine-nucleotide deletion within the S gene encoding the RBD. These viruses belong to the B.1.1.7 lineage, which had already acquired two independent deletions in recurrent deletion region ^6^ (RDR) 1 (Δ69-70) and RDR2 (Δ144/145). Spread via human-to-human transmission is likely given the timing of sample collection, geographic proximity and the clustering of sequences within a phylogeny of contemporaneously circulating B.1.1.7 isolates (Fig. 1).

**Fig. 1.**
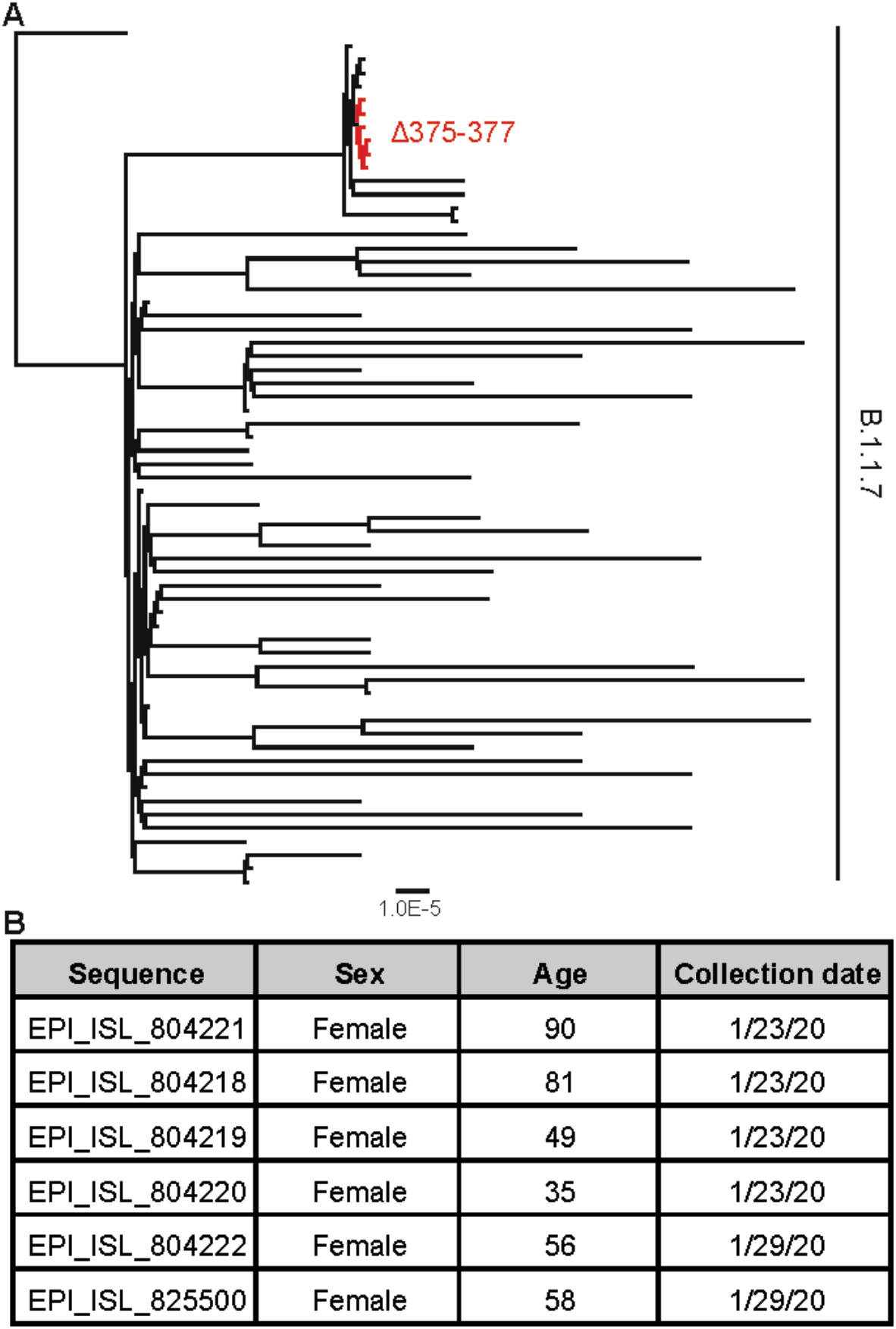
Transmission cluster within the B.1.1.7 lineage with a novel deletion in RBD. **a**. SARS-CoV-2 genome sequences with a nine-nucleotide deletion at codons 375-377 (red branches) cluster together among contemporaneously circulating B.1.1.7 isolates (black branches). The maximum likelihood phylogenetic tree is, rooted on EPI_ISL_581117 and was calculated with 10,000 bootstrap replicates. **b**. GISAID accession numbers and metadata from the six individuals.

The deletion removes residues 375-377 of the RBD. All three have been conserved for the duration of the pandemic. We examined their conservation among divergent sarbecoviruses, including SARS-CoV, bat and pangolin sequences (Fig. 2a). Among these isolates the three codons differ only by synonymous nucleotide substitutions suggesting selective pressures to preserve the identity of each amino acid. Residues 375-377 contribute to an extended surface that is broadly conserved among sarbecovirus (Fig. 2b). This site is distal from the interface between RBD and its receptor angiotensin-converting enzyme 2 (ACE2) ^7^. In the “three RBD” down state of the S glycoprotein trimer, residues 375-377 form a β-strand which is recessed and facing inward to the trimer 3-fold axis of symmetry. Sampling of “RBD-up” conformations and receptor engagement exposes this site.

**Fig. 2.**
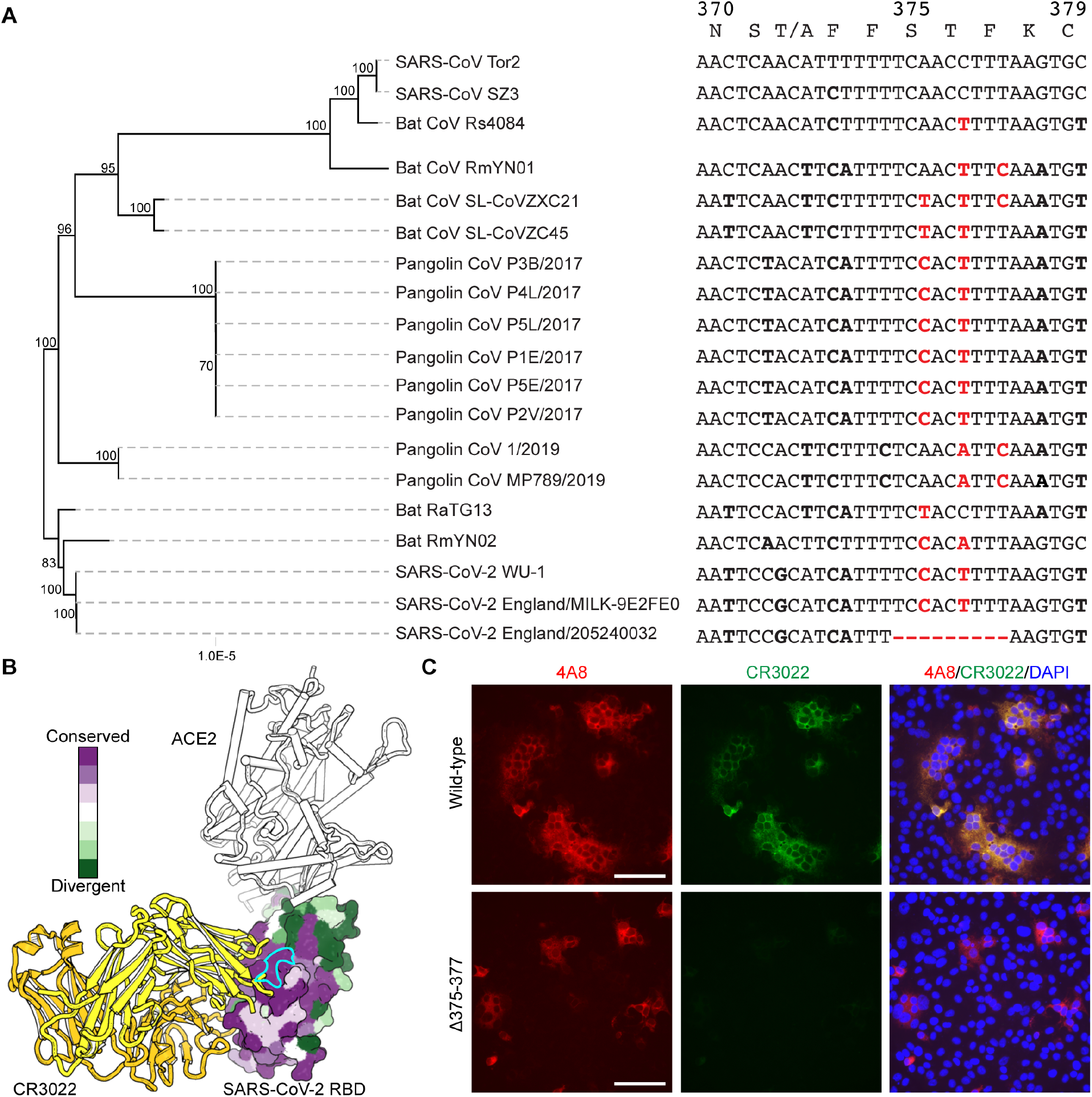
Deletion of a conserved site abolishes CR3022 binding. **a**. phylogeny of sarbecovirus genomes and nucleotide sequences of the deletion region. Differences from SARS-CoV Tor2 are shown in bold and in red for codons 375-377. SARS-CoV Tor2 specific translation and amino acid numbering is above. The maximum likelihood phylogenetic tree was calculated with 10,000 bootstrap replicates. **b**. A structure of SARS-CoV-2 bound with ACE2 (white) (PDB: 6M0J). CR3022 (6W41) (yellows) has been docked in by aligning ACE2 RBDs. The RBD surface is colored by conservation among the coronavirus isolates above using ConSurf ^28,29^ and residues 375-377 are circled in cyan. S glycoprotein distribution in Vero E6 cells at 24 h post-transfection with plasmids encoding SARS-CoV-2 S-Δ375-377 or wild-type S glycoprotein, visualized by indirect immunofluorescence in permeabilized cells. A monoclonal antibody against the N-terminal domain of SARS-CoV-2 S protein (4A8; red) detects wild-type and mutant protein. CR3022 monoclonal antibody (CR3022; green) does not detect the S-Δ375-377 mutant. Overlay images (4A8/CR3022/DAPI) depict co-localization of the antibodies; nuclei were counterstained with DAPI (blue). The scale bars represent 100 µm.

Despite its transient exposure, the conserved surface is immunogenic. A number of reported antibodies, isolated from different donors, engage this site and can bind/neutralize other sarbecoviruses. Antibody CR3022, elicited by a SARS-CoV infection during the 2003-2004 outbreak, initially defined this antibody class and their common epitope ^8-10^ (Fig. 2b). These antibodies inhibit viral replication but have limited neutralization potency in single round infection assays ^11-13^. CR3022, like others makes direct contacts with residues 375-377. We introduced Δ375-377 into an S glycoprotein expression construct and detected expression by indirect immunofluorescence using an N-terminal domain binding antibody, 4A8^14^. The formation of multinucleated, syncytia demonstrates S-Δ375-377 remains a functional membrane fusogen (Fig. 2c). However, the deletion abolishes CR3022 binding. While evolutionary conserved, this site is mutable and a single mutation event results in antibody escape.

SARS-CoV-2 only recently crossed into humans. Mounting evidence suggests that specific variants of concern have evolved some resistance to dominant humoral responses. Specifically how SARS-CoV-2 will adapt to immune pressures imposed by a human antibody repertoire is to be determined. This transmitted variant, with a deletion at an otherwise conserved site demonstrates that antigenic stability in animal species may not always extend to humans. Focused genetic surveillance has not identified additional Δ375-377-linked cases. The virus was sufficiently fit to transmit between at least five individuals and to define their viral consensus sequences. This early period of S evolution has been defined by recurrent, convergent evolution. Many defining mutations in current variants of concern are identical or functionally equivalent. The alteration of a conserved epitope by Δ375-377 not only represents an additional antigenic step in a variant of concern (B.1.1.7), but also demonstrates a capacity of this site to acquire antibody resistance rapidly.

The emergence of variants of concern and their continued evolution demonstrate that S is not as antigenically stable as initially hypothesized ^15,16^. Second generation vaccines may selectively focus antibodies to conserved epitopes, either by design or as a consequence of recalling responses from immunologic memory. The conserved site tolerates both S-Δ375-377 and mutations about its periphery ^17-20^. It may not be a suitable target for therapeutic antibodies or immune focusing immunogens.

## Acknowledgements

We gratefully acknowledge the authors from the originating laboratories and the submitting laboratories, who generated and shared via GISAID genetic sequence data on which this research is based (Table 1). We sincerely apologize to the many authors we could not cite due to limitations on references. This work was supported by The University of Pittsburgh, the Center for Vaccine Research (WPD and KRM), The Richard King Mellon Foundation, the Henry L. Hillman Foundation and the Commonwealth of Pennsylvania, Department of Community and Economic Development (WPD)

## Author contributions

L.J.R., L.R.R-M, S.N, W.P.D and K.R.M designed the experiments; L.J.R., L.R.R-M, S.N and K.R.M. performed the experiments; L.J.R., L.R.R-M, S.N, W.P.D and K.R.M. analyzed data and L.J.R., L.R.R-M, S.N, W.P.D and K.R.M wrote the manuscript.

## Methods

### Materials and Methods

#### Sequence analysis

Sequences were obtained from the publicly available GISAID database ^5^ and acknowledged in supporting Table 1. Sequence analysis was performed in Geneious (Biomatters, New Zealand). To identify deletion variants in S gene, sequences were mapped to NCBI reference sequence MN985325 (SARS-CoV-2/human/USA/WA-CDC-WA1/2020), the S gene open reading frame was extracted, remapped to reference and parsed for deletions using a search for gaps function.

All sequences were aligned in MAFFT ^21,22^ and adjusted manually for consistency. To evaluate the phylogenetic relationships between S-Δ375-377 variants and isolates from contemporaneously circulating B.1.1.7 we obtained sequences of B.1.1.7 variants from the United Kingdom from samples that were obtained from Mid-late December 2020. The first sequenced B.1.1.7 isolate hCoV-19/England/MILK-9E2FE0/2020 (EPI_ISL_581117), was included. FastTree ^23^ was used to generate a preliminary phylogeny. Branches from a node containing S-Δ375-377 variants were extracted and along with hCoV-19/England/MILK-9E2FE0/2020 (EPI_ISL_581117) were re-aligned. The final Maximum-Likelihood phylogenetic trees were calculated using RAxML ^24^ using a general time reversible model with optimization of substitution rates (GTR GAMMA setting), starting with a completely random tree, using rapid Bootstrapping and search for best-scoring ML tree with 10,000 bootstraps of support performed. The phylogeny of sarbecoviruses used the indicated sequences and were produced using RAxML ^24^ with the same parameters as above.

#### Cell lines

Human 293F cells were maintained at 37° Celsius with 5% CO_2_ in FreeStyle 293 Expression Medium (ThermoFisher) supplemented with penicillin and streptomycin. Vero E6 cells were maintained at 37° Celsius with 5% CO_2_ in high glucose DMEM (Invitrogen) supplemented with 1% (v/v) Glutamax (Invitrogen) and 10% (v/v) fetal bovine serum (Invitrogen).

#### Recombinant IgG expression and purification

The heavy and light chain variable domains of 4A8 ^14^ and CR3022 ^8^ were synthesized by Integrated DNA Technologies (Coralville, Iowa) and cloned into a modified human pVRC8400 expression vector encoding for full length human or mouse IgG1 heavy chains and human kappa light chains ^25^. IgGs were produced by polyethylenimine (PEI) facilitated, transient transfection of 293F cells that were maintained in FreeStyle 293 Expression Medium. Transfection complexes were prepared in Opti-MEM and added to cells. Five days post-transfection supernatants were harvested, clarified by low-speed centrifugation, adjusted to pH 5 by addition of 1 M 2-(N-morpholino)ethanesulfonic acid (MES) (pH 5.0), and incubated overnight with Pierce Protein G agarose resin (Pierce, ThermoFisher). The resin was collected in a chromatography column, washed with a column volume of 100 mM sodium chloride 20 mM (MES) (pH 5.0) and eluted in 0.1 M glycine (pH 2.5) which was immediately neutralized by 1 M TRIS(hydroxymethyl)aminomethane (pH 8). IgGs were then dialyzed against phosphate buffered saline (PBS) pH 7.4.

#### Cloning and transfection of SARS-CoV-2 spike protein deletion mutants

S-Δ375-377 was generated in HDM_SARS2_Spike_del21_D614G ^26^ a plasmid containing SARS-CoV-2 S protein lacking the 21 C-terminal amino acids. HDM_SARS2_Spike_del21_D614G was a gift from Jesse Bloom (Addgene plasmid # 158762; http://n2t.net/addgene:158762; RRID:Addgene_158762). The region containing amino acids 375-377 was excised using appropriate restriction enzymes and replaced by a synthetically generated gBlock (Integrated DNA Technologies) with amino acids 375-377 deleted. The Gibson Assembly reaction, transformations, clone screening, plasmid DNA preparation and transfections were performed as described previously ^6^

#### Indirect immunofluorescence

Indirect immunofluorescence was performed as previously reported ^27^. Cells were transfected with the SARS-CoV-2 S-Δ375-377 protein deletion mutant and controls. Primary monoclonal antibodies were 4A8 (mouse Fc; 1 µg/ml) and CR3022 (human Fc; 4 µg/ml), and secondary antibodies were goat anti-mouse Alexa Fluor-568, Invitrogen, and goat anti-human Alexa Fluor-488, Invitrogen, both used at 1:400 dilution.

#### Structure visualization

Structural figures were rendered in Pymol (The PyMOL Molecular Graphics System, Version 2.0 Schrödinger, LLC).

## References

1 Baden, L. R. et al. Efficacy and Safety of the mRNA-1273 SARS-CoV-2 Vaccine. N Engl J Med 384, 403–416, doi:10.1056/NEJMoa2035389 (2021).

2 Logunov, D. Y. et al. Safety and efficacy of an rAd26 and rAd5 vector-based heterologous prime-boost COVID-19 vaccine: an interim analysis of a randomised controlled phase 3 trial in Russia. Lancet 397, 671–681, doi:10.1016/S0140-6736(21)00234-8 (2021).

3 Polack, F. P. et al. Safety and Efficacy of the BNT162b2 mRNA Covid-19 Vaccine. N Engl J Med 383, 2603–2615, doi:10.1056/NEJMoa2034577 (2020).

4 Voysey, M. et al. Safety and efficacy of the ChAdOx1 nCoV-19 vaccine (AZD1222) against SARS-CoV-2: an interim analysis of four randomised controlled trials in Brazil, South Africa, and the UK. Lancet 397, 99–111, doi:10.1016/S0140-6736(20)32661-1 (2021).

5 Shu, Y. & McCauley, J. GISAID: Global initiative on sharing all influenza data - from vision to reality. Euro Surveill 22, doi:10.2807/1560-7917.ES.2017.22.13.30494 (2017).

6 McCarthy, K. R. et al. Recurrent deletions in the SARS-CoV-2 spike glycoprotein drive antibody escape. Science, eabf6950, doi:10.1126/science.abf6950 (2021).

7 Lan, J. et al. Structure of the SARS-CoV-2 spike receptor-binding domain bound to the ACE2 receptor. Nature 581, 215–220, doi:10.1038/s41586-020-2180-5 (2020).

8 er Meulen, J. et al. Human monoclonal antibody combination against SARS coronavirus: synergy and coverage of escape mutants. PLoS Med 3, e237, doi:10.1371/journal.pmed.0030237 (2006).

9 Tian, X. et al. Potent binding of 2019 novel coronavirus spike protein by a SARS coronavirus-specific human monoclonal antibody. Emerg Microbes Infect 9, 382–385, doi:10.1080/22221751.2020.1729069 (2020).

10 Yuan, M. et al. A highly conserved cryptic epitope in the receptor binding domains of SARS-CoV-2 and SARS-CoV. Science 368, 630–633, doi:10.1126/science.abb7269 (2020).

11 Huo, J. et al. Neutralization of SARS-CoV-2 by Destruction of the Prefusion Spike. Cell Host Microbe 28, 445–454 e446, doi:10.1016/j.chom.2020.06.010 (2020).

12 Wrobel, A. G. et al. Antibody-mediated disruption of the SARS-CoV-2 spike glycoprotein. Nat Commun 11, 5337, doi:10.1038/s41467-020-19146-5 (2020).

13 Finkelstein, M. T. et al. Structural Analysis of Neutralizing Epitopes of the SARS-CoV-2 Spike to Guide Therapy and Vaccine Design Strategies. Viruses 13, doi:10.3390/v13010134 (2021).

14 Chi, X. et al. A neutralizing human antibody binds to the N-terminal domain of the Spike protein of SARS-CoV-2. Science 369, 650–655, doi:10.1126/science.abc6952 (2020).

15 Dearlove, B. et al. A SARS-CoV-2 vaccine candidate would likely match all currently circulating variants. Proc Natl Acad Sci U S A 117, 23652–23662, doi:10.1073/pnas.2008281117 (2020).

16 Rausch, J. W., Capoferri, A. A., Katusiime, M. G., Patro, S. C. & Kearney, M. F. Low genetic diversity may be an Achilles heel of SARS-CoV-2. Proc Natl Acad Sci U S A 117, 24614–24616, doi:10.1073/pnas.2017726117 (2020).

17 Greaney, A. J. et al. Comprehensive mapping of mutations to the SARS-CoV-2 receptor-binding domain that affect recognition by polyclonal human serum antibodies. bioRxiv, 2020.2012.2031.425021, doi:10.1101/2020.12.31.425021 (2021).

18 Greaney, A. J. et al. Complete Mapping of Mutations to the SARS-CoV-2 Spike Receptor-Binding Domain that Escape Antibody Recognition. Cell Host Microbe 29, 44–57 e49, doi:10.1016/j.chom.2020.11.007 (2021).

19 Starr, T. N. et al. Deep Mutational Scanning of SARS-CoV-2 Receptor Binding Domain Reveals Constraints on Folding and ACE2 Binding. Cell 182, 1295–1310 e1220, doi:10.1016/j.cell.2020.08.012 (2020).

20 Liu, Z. et al. Identification of SARS-CoV-2 spike mutations that attenuate monoclonal and serum antibody neutralization. Cell Host Microbe, doi:10.1016/j.chom.2021.01.014 (2021).

21 Katoh, K., Misawa, K., Kuma, K. & Miyata, T. MAFFT: a novel method for rapid multiple sequence alignment based on fast Fourier transform. Nucleic Acids Res 30, 3059–3066, doi:10.1093/nar/gkf436 (2002).

22 Katoh, K. & Standley, D. M. MAFFT multiple sequence alignment software version 7: improvements in performance and usability. Mol Biol Evol 30, 772–780, doi:10.1093/molbev/mst010 (2013).

23 Price, M. N., Dehal, P. S. & Arkin, A. P. FastTree: computing large minimum evolution trees with profiles instead of a distance matrix. Mol Biol Evol 26, 1641–1650, doi:10.1093/molbev/msp077 (2009).

24 Stamatakis, A. RAxML version 8: a tool for phylogenetic analysis and post-analysis of large phylogenies. Bioinformatics 30, 1312–1313, doi:10.1093/bioinformatics/btu033 (2014).

25 Watanabe, A. et al. Antibodies to a Conserved Influenza Head Interface Epitope Protect by an IgG Subtype-Dependent Mechanism. Cell 177, 1124–1135 e1116, doi:10.1016/j.cell.2019.03.048 (2019).

26 Crawford, K. H. D. et al. Protocol and Reagents for Pseudotyping Lentiviral Particles with SARS-CoV-2 Spike Protein for Neutralization Assays. Viruses 12, doi:10.3390/v12050513 (2020).

27 Klimstra, W. B. et al. SARS-CoV-2 growth, furin-cleavage-site adaptation and neutralization using serum from acutely infected hospitalized COVID-19 patients. J Gen Virol, doi:10.1099/jgv.0.001481 (2020).

28 Ashkenazy, H. et al. ConSurf 2016: an improved methodology to estimate and visualize evolutionary conservation in macromolecules. Nucleic Acids Res 44, W344–350, doi:10.1093/nar/gkw408 (2016).

29 Celniker, G. et al. ConSurf: Using Evolutionary Data to Raise Testable Hypotheses about Protein Function. Israel Journal of Chemistry 53, 199-206, doi:https://doi.org/10.1002/ijch.201200096 (2013).

